# P300 Parameters Over the Lifespan: Validating Target Ranges from an In-Clinic Platform

**DOI:** 10.1101/2021.10.25.465715

**Authors:** D. S. Oakley, K. Fosse, E. Gerwick, D. Joffe, D. A. Oakley, A. Prather, F Arese Lucini, F. X. Palermo

## Abstract

**Background:** Evoked potential (ERP) markers extracted from an EEG exam can provide novel sources of information regarding brain function allowing changes in cognition, from conditions such as concussion or unhealthy aging, to be measured and tracked over time. Age-related targets can provide useful guides for practitioners, trainers, and patients seeking to optimize and track brain function over time.

Objective 1: To compare age-related changes in auditory P300 metrics of latencies and amplitudes collected in the course of routine clinical evaluations with published trends from research.
Objective 2: To establish within-person test-retest variances for these P300 metrics.
Objective 3: To validate a method for presenting age-stratified target ranges of P300 metrics.

**Participants:** One thousand seven-hundred and seventy-eight reference subjects aged 13-90.

**Methods:** Audio P300 was measured as part of a health screening exam for studies through Colorado University, Children’s Hospital Colorado, Boone Heart Institute, WAVi Co., and various clinics alongside other clinical evaluations.

**Results:** The age-related trends in both the P300 latency and amplitude measured in clinic match the age-related trends from previous research. In particular, the 2 control group endpoints at 20 and 90 years match the linear slopes predicted for age-related decline from early studies, and the inclusion of all ages produces a maturation prediction of a recent meta analysis where metrics peak near 20 years of age. Large between-person variances are observed across all studies but within-person variances remain consistent with previous studies.

**Conclusion:** In-clinic measures of P300 latency and amplitude corroborate the age-related trends of published research taken over the last several decades. Large between-person variance remains, leaving P300 best suited for within-person comparison.

## Introduction

Event-related potentials (ERP) are a measurement of the electroencephalogram (EEG) signal time-locked to the onset of a given stimulus and consist of different components labeled by their polarity (P for positive or N for negative) and their time of occurrence after the stimulus in milliseconds (e.g. P300).

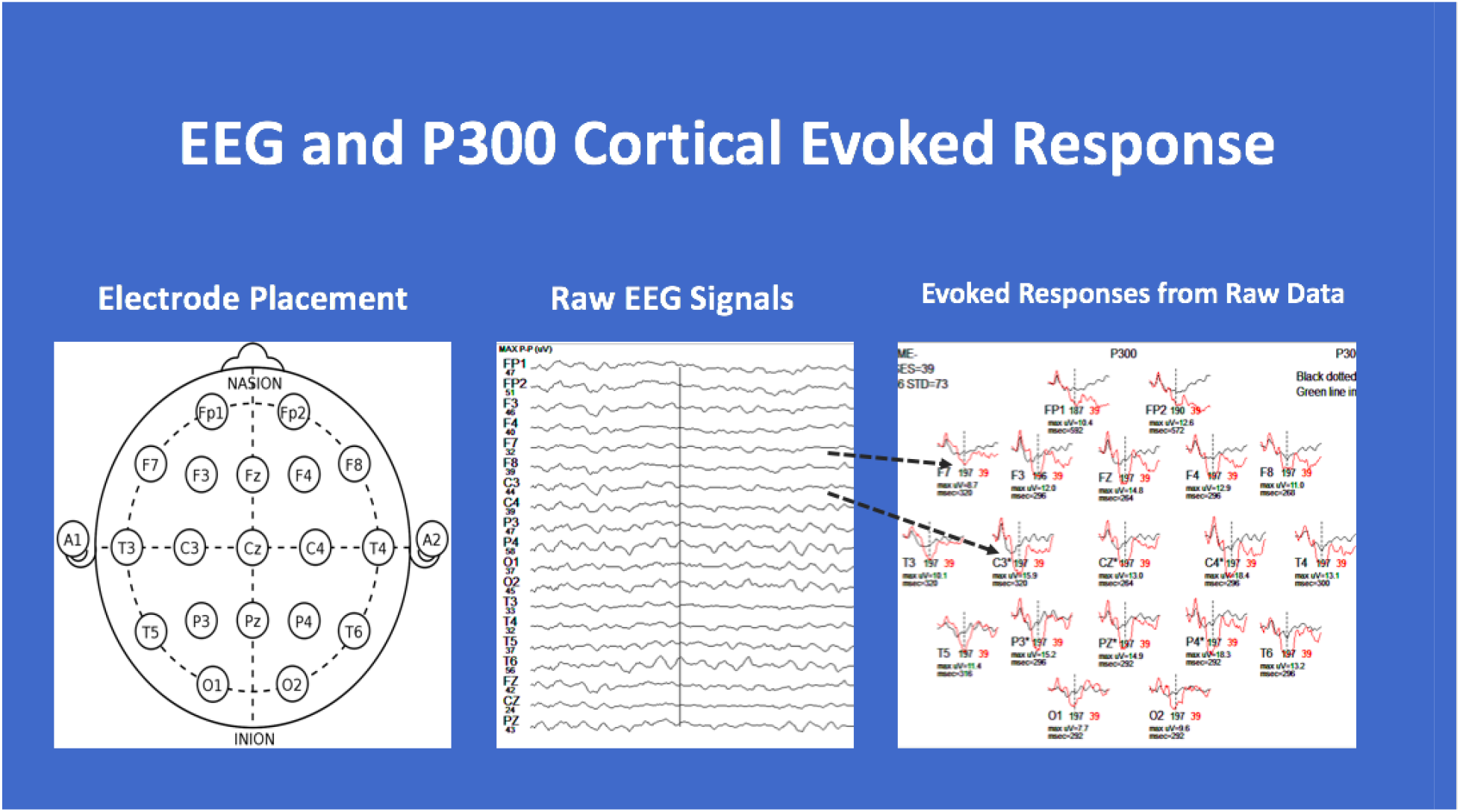

ERP waveforms are extracted by taking averages of EEG signal, according to scalp location, over a specified time period after delivery of the stimulus (Figure 1). On the far right plot of Figure 1, the red lines represent the averaged voltage measured during the rare tone, black the common, and the vertical dashed line represents 300ms post stimulus where the P300 trough is typically found (a positive voltage displayed in the downward directions as per convention).

The oddball P300 protocol, one of the most widely studied, utilizes measurements of amplitude and latency to assess the brain’s cognitive ability to recognize the odd tone as different. P300 amplitude is thought to be proportional to the amount of attentional resources devoted to a given task where P300 latency (the delay between stimulus delivery and recognition of the oddball tone as different) is a measure of stimulus classification speed. ^1^ An increase in the P300 peak latency, and/or a decrease in the P300 peak amplitude are observed in various conditions accompanied by a decline in cognitive function, including aging, dementia, mild traumatic brain injury (mTBI), substance abuse, depressive disorders, cardio health, pre-Alzheimer’s, responsivity to pharmaceuticals, trauma, and many others.^2 3 4 5^ *Decreased* latencies and *increased* amplitudes, on the other hand, have been shown to correlate with *better* cardiovascular health and more *successful* cognitive aging. ^6 7^

While significant differences are often seen between these various groups and controls, person-to-person variances may make it difficult to draw conclusions about a single individual from a single test. ^8^ Hence, these techniques are still largely utilized only in research. The variance within a person, however, is small enough to allow for these techniques to be used for longitudinal tracking and thereby as cognitive-screening tools.^9 10^ For example, recent studies have shown that if ERP’s are utilized in baseline-screening the resulting changes in P300 are highly sensitive to concussion, improved cardio health, and improved cognition after pharmaceutical intervention. ^6, 9, 11^

The other studies correlating P300 changes to cognitive impairment underscore the important information P300 can add if baseline values are established and measured during routine physical evaluations. WAVi has created user interfaces that allows for efficient longitudinal tracking that is being used in research that enables studies to take place in real-life clinical settings. The purpose of this paper is to validate this in-clinic data against published trends—trends compiled over the last 4 decades in numerous studies. In particular, we will compare methods for displaying exam results against trends. To that end the objective of this paper is to (1) To compare age-related changes in auditory P300 metrics of latencies and amplitudes collected in the course of routine clinical evaluations with published trends from research. (2) To establish within-person test-retest variances for these P300 metrics collected (3) To validate a method for presenting age-stratified target ranges of P300 metrics.

### P300 as a Brain Activity Measure Over the Lifespan

The oddball audio P300 protocol is well suited for in-clinic measurements because it is readily standardizable, has been well studied, and can be implemented on a large scale.^9^ The P300 response has also been extensively studied across the lifespan, which include amplitude decreases and latency increases with age fit with linear slopes and a more recent meta analysis showing that latency and amplitude follow a maturational path from childhood to adolescence and then where degenerative effects begin. ^2, 3, 7^ Here it was hypothesized that the P300 latency indexes neural speed or brain efficiency and the P300 amplitude index neural power or cognitive resources, which increase with maturation and then begin to decrease at older ages. We will compare both of these models to our data.

## Methods

### Subjects

While the subjects for this study were comprised of 1778 subjects from previous or ongoing studies, it is not intended that they represent a normal control for a general population, rather it is intended to provide a target reference. One of the goals of this study is to compare data collected during routine clinical evaluations with historical research to test the validity of large-scale screening. This study does have 3 control groups (13-16 years of age, 17-23, and 81-90) and these will anchor the resulting age-matched curves as discussed below. It may be the case that these controls perform differently from those found in a normal population, where 2 of these control groups were taken from elite club, High School, or NCAA athletic teams while those in the oldest age range were volunteers living independently, still interested in brain science, and still interested in their brain performance. Each group will be discussed individually, but because of the suspected other-than-normal performance, we will focus on age trends and refer to this reference group as a target reference, with end points as discussed, rather than a normal reference. Also, in keeping with the focus of this paper which is to establish age-related reference targets and compare these to literature trends, potential male-female differences are not investigated here.

All studies were approved by appropriate IRB’s and written informed consent was obtained from the participants before study intake.

#### Ages 8-12

48 subjects aged 8-12 were taken from three previous studies: a study that followed athletes over the course of their sports seasons in Texas and Washington, control subjects measured as part of a beta test to explore the outcome of an educational/wellbeing intervention program in an economically-challenged school, ^12^ and wards accompanying WAVi study volunteers discussed below.

#### Ages 13-16

This control group comprises 83 subjects from a previous study following 94 athletes aged 13-16 over the course of their sports seasons and at 4 different sites. These subjects are participants in youth soccer and youth basketball representing all players from single teams. ^13^

In these previous studies, these subjects were controls against which pre-contact, postconcussion and return-to-play groups could be compared. Here P300 voltages, along with reaction time and Trail Making measurements, were assessed during the course of other precontact clinical evaluations administered by sports medicine staff. To follow the objectives of this study (as well as the above-mentioned studies) which involves real clinical settings, and because the primary marker being studied is nonspecific, our exclusion criteria are minimal. The “control” group, therefore, is a reference group taken from all players participating on these teams with no exclusions, except that those who had lower than 80% yield on the audio P300 protocol that were excluded due to artifact. Of these 83 subjects, 66 returned and completed a valid post-season second test which will be used to discuss test-retest variability.

#### Ages 17-23

This second control group is taken from a previous study that followed 364 athletes aged 17-23 over the course of up to 4 sports seasons and at 5 different sites. These subjects are participants in NCAA Div. 1 men’s football (172 players, representing all players from a single team), woman’s soccer (29 NCAA Div. 1, representing all players from a single team), men’s high school football (142 players, representing all seniors from a single team), and semipro men’s ice hockey (20 players, representing all players from a single team). ^9^

In these previous studies, these subjects were controls against which pre-contact, postconcussion and return-to-play groups could be compared. Here P300 voltages, along with reaction time and Trail Making measurements, were assessed during the course of other precontact clinical evaluations administered by sports medicine staff on 356 of the 364 players tracked. To follow the objectives of this study (as well as the above-mentioned studies) which involves real clinical settings, and because the primary marker being studied is nonspecific, our exclusion criteria are minimal. The “control” group, therefore, is a reference group taken from all players participating on these teams and exclusions are limited to the players who fell asleep during the first-year test and passing the artifact criteria discussed above, leaving a total of 304 players comprising the baseline reference group of Table I. Of these subjects, 70 returned injury free to complete a valid second test, which will be used to discuss test-retest variability for P300 amplitude, with a subset of 38 of these to test P300 latency variation (because of a change in protocol as noted in the study).^9^

**Table I:** Profile of Assessments.

| **Table I: Profile of Assessments.** |  |  |
| --- | --- | --- |
| **Age (yrs)** | **# Assessments included** | **Subject Profile** |
| 8-12 | 48 | Youth Sports and Educational studies: *initial assessment upon enrollment into study.* |
| 13-16 | 83 | Youth Sports study: *initial pre-contact assessment upon enrollment into study.* |
| 17-23 | 304 | High School and University Soccer, Hockey, and US Football study: *initial pre-contact assessment upon enrollment into study.* |
| 24-30 | 128 | Preventative Cardiology and Healthy Aging studies: *initial pre-intervention assessment upon enrollment into study.* |
| 31-40 | 204 | Preventative Cardiology and Healthy Aging studies: *initial pre-intervention assessment upon enrollment into study.* |
| 41-50 | 297 | Preventative Cardiology and Healthy Aging studies: *initial pre-intervention assessment upon enrollment into study.* |
| 51-60 | 377 | Preventative Cardiology and Healthy Aging studies: *initial pre-intervention assessment upon enrollment into study.* |
| 61-70 | 231 | Preventative Cardiology and Healthy Aging studies: *initial pre-intervention assessment upon enrollment into study.* |
| 71-80 | 56 | Preventative Cardiology and Healthy Aging studies: *initial pre-intervention assessment upon enrollment into study.* |
| 81-90 | 42 | Healthy Aging study: *initial assessment upon enrollment into study.* |
| 24-85 | 67 (test-retest) | Eyes-closed Resting and Audio P300 study: *67 test-retest assessments on 8 subjects testing audio P300 as an eyes-closed qEEG test.* |

#### Ages 24-30

128 individuals aged 24-30 were measured in clinic at baseline where some were to be tracked over the course of various interventions. Subjects include patients who visited the Boone Heart Institute Colorado for a combined preventative cardiology and EEG/ERP evaluation from June 2014 through June 2017. Only first-time patients receiving an initial evaluation were included in the sample, which was also used for a preventative cardiology study. ^6^ Because this is a target reference study, the exclusion criteria are minimal, the criteria being those who were taking beta-blockers or psychiatric medication and those who had lower than 80% yield on evoked responses due to artifact.

Also included were subjects from Natural Bio Health (NBH) Texas for a first-time preventative wellness exam, evaluated from 2017 through 2018; and a random sampling of subjects measured for demonstration purposes at 5 medical conferences.

The remaining subjects were volunteers who were known to or associated with the study team and wanted to become proactive in their brain health. In general these reference subjects were well educated and wanted to use WAVi to compare pre-intervention to post-interventions where interventions typically included some form of lifestyle change. To follow the objectives of this study, which involves real clinical settings, and because the primary marker being studied is nonspecific, our exclusion criteria are minimal and all volunteers in this age group were analyzed for the purposes of this study.

#### Ages 31-40

204 individuals aged 31-40 were tracked over the course of various interventions and the above-mentioned clinics, conferences, and volunteers. Of these, 56 were selected to be used as controls for other studies who had no history of trauma and were not on any medications. We report these subjects here.

#### Ages 41-50

297 individuals aged 41-50 were tracked over the course of various interventions and the above-mentioned clinics, conferences, and volunteers. Of these, 56 were selected to be used as controls for other studies who had no history of trauma and were not on any medications.^14^ We report these subjects here.

#### Ages 51-60

377 individuals aged 31-40 were tracked over the course of various interventions and the above-mentioned clinics, conferences, and volunteers.

#### Ages 61-70

231 individuals aged 31-40 were tracked over the course of various interventions and the above-mentioned clinics, conferences, and volunteers.

#### Ages 71-80

56 individuals aged 31-40 were tracked over the course of various interventions and the above-mentioned clinics, conferences, and volunteers.

#### Ages 81-90

Our third control group comprises 42 people taken as volunteers, discussed above. This group was living independently, had not been diagnosed with dementia, and were by definition a population who had experienced what could be called successful cognitive aging. They provide an end point against the 20-year old athletes for our target reference.

#### Ages 24-61 Test-Retest Group

This group comprises 67 tests from 8 volunteers to study test-retest for various ERP and qEEG parameters during eyes-closed audio P300.

### EEG acquisition and preprocessing

After completing a short 1-page intake form, participants were given an EEG test that included the oddball audio P300 component. Reaction times were also measured by asking the participant to click the track pad or mouse upon hearing the oddball tone.

The EEG was recorded using the WAVi ™ Research Platform (WAVi Research, Boulder, CO, USA) sampled at 250 Hz and bandpass filtered between 0.5-30 Hz. The electrodes were placed according to the International 10–20 system using caps with 19 tin electrodes (both with the WAVi Headset and Electro-cap International Inc., Eaton, Ohio, USA). Linked reference electrodes were placed at the earlobes.

The test administrators were instructed to keep the electrode impedances below 20kOhm for EEG locations and below 10kOhm for the ground-to-ear locations where possible. These targets are well below the 1 GOhm input impedance of the WAVi amplifiers, are practical regarding preparation time in the sports setting (including certain “sport-related” hairstyles) and produced sufficient yield.^15^

To be consistent with the goal of testing a simplified platform, a continuous 4 minute 2-tone audio oddball eyes-closed P300 protocol was used. Here, 200 common tones (1000 Hz) and 40 rare tones (2777 Hz) were delivered in random order over the span of 4 minutes, creating a 0.95s interstimulus interval with a 50 ms tone length. The tones were delivered using Skullcandy™ over the ear headphones, at 65 dB.

### EEG extraction

Individual ERPs were extracted using WAVi Scan 1.0 software (the results of which are intended to create the target ranges for WAVi Desktop and WAVi Scan software). There are different methods of extracting ERP values. The WAVi Brain Assessment reports P300 components by identifying the positive extremum in the latency range of 240–500ms. The depth (P300V) is then extracted for each electrode site from the mean amplitude of all 4 0 stimuli in this 240-500ms time window relative to the first 16 ms post stimulus, baseline corrected using the 100ms pre-stimulus period. The P300 latency (P300T) is the calculated delay of the maximum P300 value measured from the time of stimulus delivery for each electrode site.

P300 measurements are often reported from specific sites such as Cz and Pz, or from the average of the six central-parietal sites (C3, Cz, C4, P3, Pz, P4). In a slight deviation from the published trends against which we are comparing, we report the lowest recorded latency and highest amplitude from the six central-parietal sites as this was found to have the lowest test-retest variance on an individual. ^9^

All trials include automatic artifact rejection that exclude any errors from averaging, where noise from EEG data with higher than acceptable amplitudes and excessive band frequency activities in the standard EEG bands (Delta, Theta, Alpha, and Beta) were excluded on an individual channel basis. The artifacting was manually inspected to confirm proper noise extraction. Only tests with greater than an 80% yield were included in these calculations.

It is important to note that when comparisons are made between P300 studies using different EEG systems (including those using previous WAVi technology) it should be noted that software filter settings can affect the overall values of P300 amplitude, and this is something often overlooked in meta-analysis literature. Two previous studies using the WAVi software for analysis, ^6, 9^ for example, used pre-WAVi 9.7.5 software with filter settings that, while consistent, did not utilize the DC-drift correction settings of later versions aimed at increasing yield in the face of differing patient headset electrode preparation experienced in real-life clinical settings. While these new filters had no effect on the overall trend, they did cause an overall 9% decrease in P300 amplitude from previous studies using the WAVi software.

## Results

### P300 Age Trends

WAVi data for P300 latency and amplitude are shown in Figures 2 and 3, and Table II. Notice the large person-to-person variance as discussed above, highlighting the need for individual baseline tracking. The black lines are the predicted linear fit of previous work, where the voltage was predicted to drop 0.1uV/yr and the latency was predicted to increase by 0.8ms/yr.^3^ These lines were normalized to the data from 20-year old athletes, in keeping with the philosophy that we are looking for a target reference range for performance comparison. Here the overall amplitude is 1uV higher than reported in meta-analyses and latency 26 ms shorter, ^7^ most-likely due to the fact that we report the highest amplitude/lowest latency in the C-P region rather than the common method of just reporting the values at Cz or Pz or an average. Anchoring the line to the 20-year old age range show shows surprising linear fits between 20 and 85 years old—surprising in part because of the number of years between this study and the previous study predicting the linear fit. These figures also compare another prediction from the meta-analysis fit suggesting modification from linear on account of maturation stages (dotted lines). ^7^ These plots were also normalized against the 20-year old athletes and show a better fit than the linear. Although we do not show data for ages younger than 8, maturation effects are also seen to emerge in amplitude declines and latency increases (delays). The best polynomial fits to these data are shown in the broad blue line and match the maturation prediction. Here the mid-age ranges show lower than predicted amplitudes and shorter latencies. It’s not clear if this is a result of data collection, which was largely from real-life clinical settings, or this represents a trend. As the goal of this work is to establish targets against which longitudinal tracking can take place, we will leave that question for future studies.

**Table II:** P300 Results.

| **Table II: P300 Results.** |  |  |  |
| --- | --- | --- | --- |
| **Age** | ***n*** | ***P300V(SD)***<br>***uV*** | ***P300T(SD)***<br>***ms*** |
| 10-12 yrs | 48 | 13.3(7.9) | 297(35) |
| 13-16 yrs | 83 | 16.9(6.6) | 294(33) |
| 17-23 yrs | 304 | 16.3(6.5) | 293(32) |
| 24-30 yrs | 128 | 12.7(5.9) | 281(32) |
| 31-40 yrs | 56 | 12.7(5.5) | 290(35) |
| 41-50 yrs | 56 | 13.0(5.2) | 289(33) |
| 51-60 yrs | 377 | 10.8(5.4) | 307(41) |
| 61-70 yrs | 231 | 10.1(5.3) | 320(47) |
| 71-80 yrs | 56 | 8.9(3.9) | 356(62) |
| 81-90 yrs | 42 | 9.5(5.0) | 362(60) |
| *Combined* | *1381* | *12.4(5.7)* | *309(42)* |

A note on variance: It is common practice in medicine to quote “normal ranges” (middle 68 percent of people in a Gaussian distribution) in order to provide context for both the clinician and client. A Gaussian distribution cannot always be assumed, however, and for P300T we have found a variance of +-30 (more stringent than Table II) provides a more useful target range.

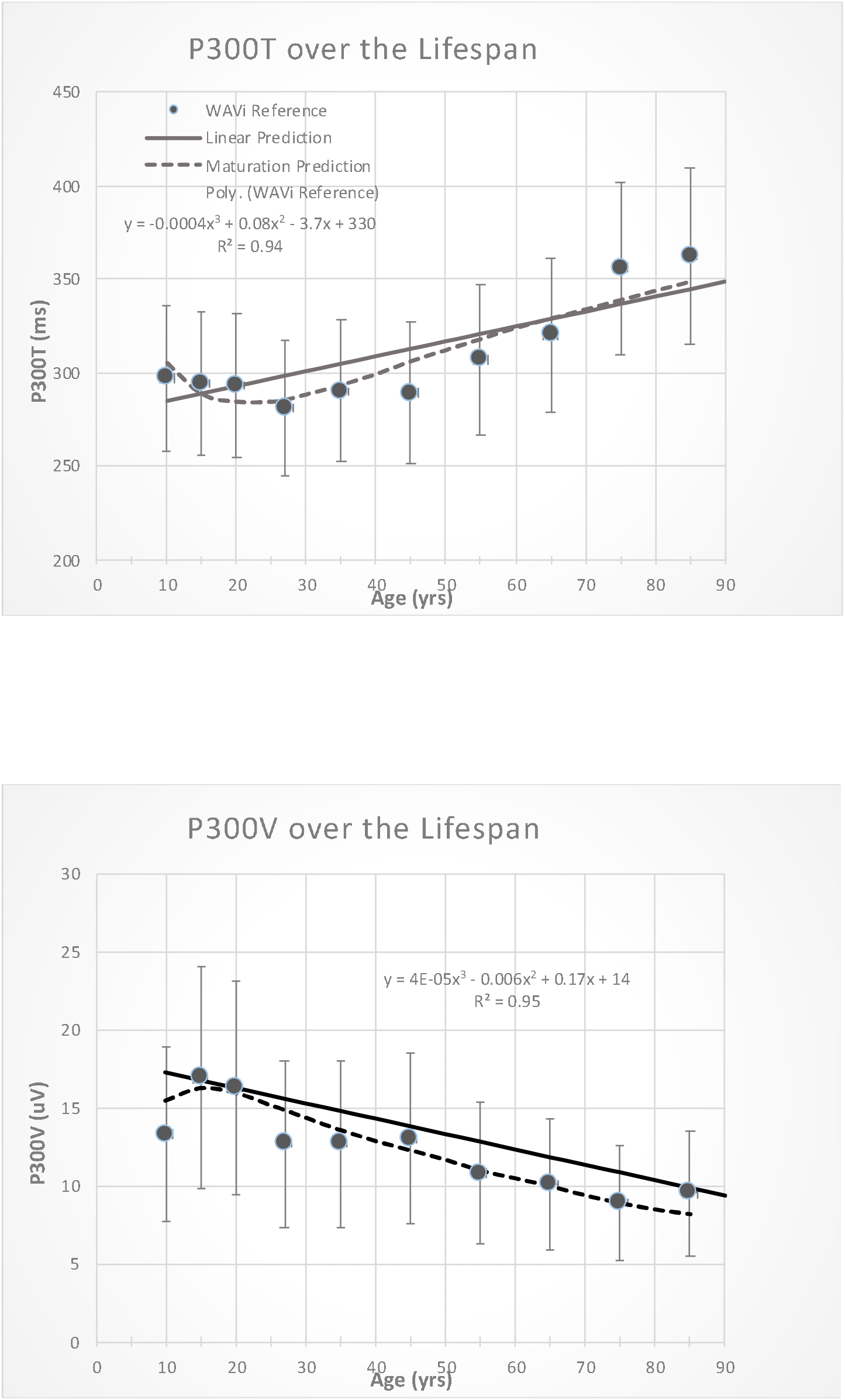

### P300 Test-Retest Variability

We can conceptualize variance in P300 metrics as a combination of trait (stable characteristics of a person), of state (representing aspects of a subjects psychological or physiologic state), and of other stimuli (such as noise in the room). To tease out the state (i.e. is there a concussion?) and for longitudinal tracking it’s therefore important to determine the within-person variance in routine clinical settings. As discussed above, while Gaussian distributions cannot be assumed for these variations, particularly with changing states, it is important to establish the top and/or bottom-performing 16% to provide clinical context.

The expected intrapersonal variance in latency and amplitude from these data is shown in Table III. Here we see previous work reporting a latency variance of +- 10 and +- 12 ms for the 13-16 and 17-23 year age groups respectively, over the course of up to 2 years between measurements. This is confirmed here in the test-retest data set covering an age range of 24-85 years where 84% of the subjects fell below +-11ms in change over the course of 1 month to 2 years. For personal variation in amplitude, previous work cited variances of +- 2uV for the 13-16 *and* 17-23 year age groups also over the course of up to 2 years. This variance is confirmed here in the test-retest data set covering an age range of 24-85 years where 84% of the subjects fell below +- 2uV in change over the course of 1 month to 2 years. These numbers provide a reference from which longitudinal P300 changes can be studied, either after an event such as concussion, an intervention, or in order to monitor healthy aging.

**Table III:**
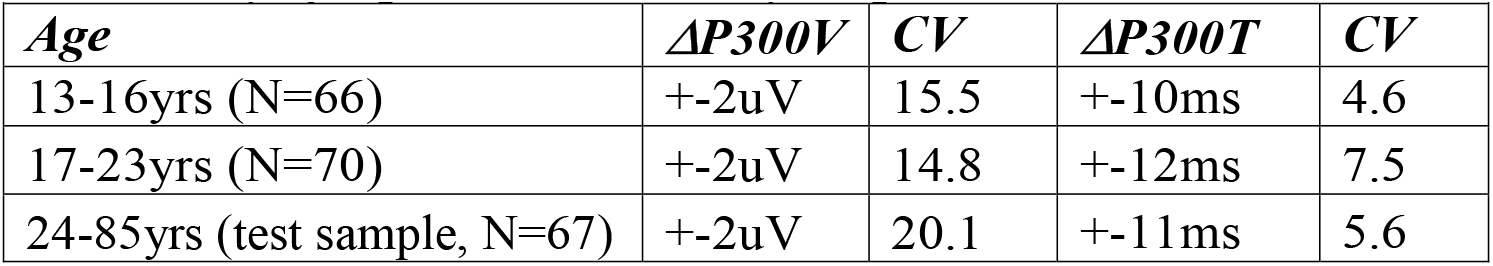
Expected longitudinal change of the P300 amplitude and latency of a person over a 0-2 year period.

Also shown in Table III are coefficients of variation (CV), calculated as 100*SD/Average for each person and then averaged over each group. This provides context to compare P300 to other methods of physiologic assay, say for example HDL measures (CV~24) and cholesterol (CV~14). ^9^ Table III CV values (comparable to a previous study of CV=5.6 for P300T and lower than the reported 31.7 for P300V) are lower than many other clinical assays, suggesting the clinical utility of longitudinal tracking. ^16^ While both P300T and P300V are presumed a combination of state and trait, the higher CV for P300V may suggest more of state or acute influences than P300T, with its lower CV suggesting more trait or chronic influences as noted in a study of heart health and P300. ^7^

### Blink Artifact P300 Test-Retest Variability

On a final note, we discuss the possibility of muscle artifact resulting from blinks that are synchronized with the delivery of the oddball tone, for example when a mouse is pressed. This situation can be more common in patients with dementia or other conditions associated with involuntary movement. Table IV shows that the muscle-related noise from eye blinks does not discernably affect the P300 signal in the C-P region. In this table we compare 8 patients with severe synchronized blinking where we analyze the P300 when allowing the blinks and when we exclude those P300 presentations where eye-movement is manually artifacted out. The small differences can be attributed to the lower yield in the manually-artifacted P300 segments.

**Table IV:** P300 amplitude and latency in the C-P region with and without blink artifact.

|  |  |  |
| --- | --- | --- |
| ***Metric*** | ***Synchronized Blinking*** | ***No Blinking*** |
| ***Largest C-P P300V (SD)*** | 15.2(6.4)uV | 15.1(6.9)uV |
| ***Average C-P P300V (SD)*** | 12.2(5.7)uV | 11.7(5.7)uV |
| ***Largest C-P P300T (SD)*** | 269(24) ms | 279(41) ms |
| ***Average C-P P300T (SD)*** | 286(36) ms | 288(44) ms |

While blinking may not have a great effect on the auditory P300 metrics in the C-P regions, it clearly influences the frontal regions. Since studies have associated conditions such as early Parkinson’s and aging to changes in the frontal P300,^17 18^ the use of these regions would require more careful artifacting, such as traditional manual inspection.

## Discussion

Data collected during routine clinical evaluations produce trends that compare to both the linear P300 decline prediction and the maturation prediction of previous studies, with a maturation prediction with maximum performance around 20 years of age providing the best fit. A target reference is a useful tool to compare trends with end points of high functioning people on both ends of the age range.

While the between-person variance can make limit the utility of these trends in single tests, the within-person test-retest variance of +-11ms for latency and +-2uV for amplitude allow for useful longitudinal tracking. Finally, care must be made when using the P300 for diagnostic purposes because these metrics are non-specific and can be affected by numerous conditions.

## Conclusion

In-clinic measures of P300 latency and amplitude corroborate the age-related trends of published research taken over the last several decades. Large between-person variance remains, leaving P300 best suited for within-person comparison where variances are sufficiently low to allow for meaningful longitudinal tracking.

## Notes

### Competing Interest Statement

D. S. Oakley, a K. Fosse, a E. Gerwick, a D. Joffe,a D. A. Oakley, a A. Prather, a F Arese Lucini are paid consultants and employees of WAVi Co, the company who's device is being validated.

